# LINE-1 Retrotransposon expression in cancerous, epithelial and neuronal cells revealed by 5’ single-cell RNA-Seq

**DOI:** 10.1101/2021.01.19.427347

**Authors:** Wilson McKerrow, Nicole Doudican, Nicholas Frazzette, Shane A. Evans, Azucena Rocha, Larisa Kagermazova, John M. Sedivy, Nicola Neretti, John Carucci, Jef D. Boeke, David Fenyö

**Affiliations:** Institute for Systems Genetics, NYU Langone Health, New York, NY, USA; Department of Biochemistry and Molecular Pharmacology, NYU Langone Health, New York, NY, USA; Ronald O. Perelman Department of Dermatology, NYU Langone Health, New York, NY, USA; Center for Computational Molecular Biology, Brown University, Providence, RI, USA; Department of Molecular Biology, Cell Biology, and Biochemistry, Brown University, Providence, RI, USA; Center on the Biology of Aging, Brown University, Providence, RI, USA; Department of Biomedical Engineering, NYU Tandon School of Engineering, Brooklyn NY 11201

## Abstract

LINE-1 retrotransposons are sequences capable of copying themselves to new genomic loci via an RNA intermediate. New studies implicate LINE-1 in a range of diseases, especially in the context of aging, but without an accurate understanding of where and when LINE-1 is expressed, a full accounting of its role in health and disease is not possible. We therefore developed a method – 5’ scL1seq – that makes use of a widely available library preparation method (10x Genomics 5’ single cell RNA-seq) to measure LINE-1 expression in tens of thousands of single cells. We recapitulated the known pattern of LINE-1 expression in tumors – present in cancer cells, absent from immune cells – and identified hitherto undescribed LINE-1 expression in human epithelial cells and mouse hippocampal neurons. In both cases, we saw a modest increase with age, supporting recent research connecting LINE-1 to age related diseases.

## Introduction

In the human genome, LINE-1 is the only family of retrotransposons – sequences capable of copying themselves to new genomic loci via an RNA intermediate – that remains active and autonomous^1^. LINE-1 element expression is driven by a promoter that lies within its 5’UTR sequence. It is hence an unusual promoter in that it contains 100% “downsteam elements”^2^, which is necessitated by its mode of retrotransposition that requires it to “take its promoter with it”. Once expressed and translated, LINE-1 mRNAs complex with their two potentially cis-acting^3^ protein products, the RNA binding ORF1p^4^ and the endonuclease/reverse transcriptase ORF2p^5,6^, to form LINE-1 RNPs that are primarily cytoplasmic but can enter the nucleus during mitosis^7^. In the nucleus, ORF2p reverse transcribes a new LINE-1 DNA copy at a site of genomic insertion through a process called target primed reverse transcription (TPRT)^8^. In addition to its potential to insert into and disrupt protein coding genes^9–12^, LINE-1 expression can contribute to DNA damage^13,14^ and induce an interferon response^14–17^. LINE-1 is highly active in cancer, with some tumors bearing more than 100 new LINE-1 insertions^18–23^. It is also active in the germline where it generates insertions that are polymorphic in the human population^24^ and occasionally contributes to heritable genetic disorders^9^.

The extent of LINE-1 activity outside these two contexts, namely in non-tumor somatic tissues, has been a subject of vigorous debate for decades^25–31^. Recently, several studies have suggested that the epigenetic changes that occur during aging provide an opportunity for LINE-1 and other normally silenced genomic regions to become active and contribute to the aging phenotype and related diseases^16,32,33^. We therefore sought a highly sensitive method that would use standard techniques to identify LINE-1 RNA expression and to ascertain what counts as normal somatic LINE-1 expression and how that changes in disease and aging contexts.

The very high copy number of LINE-1 sequences means that methods based on simple read counts of LINE-1 RNA are confounded by the fact that the hundreds of thousands of LINE-1 copies include many that lie within cellular transcription units and thus those transcripts are not driven by the LINE-1 promoter, but instead are “passively co-transcribed”. We therefore leveraged 5’ targeted single cell RNA-seq data sequenced with reads that are at least 100 base pairs long to develop 5’ scL1seq. Because 5’ scL1seq is a straightforward method to extract information about LINE-1 expression from reads sequenced from a standard library prep (5’ targeted 10x genomics single cell RNA-seq), it can easily be deployed by researchers interested in whether LINE-1 expression is present in particular disease models or samples.

We applied this method to publicly available data from 15 healthy somatic tissues, and found that LINE-1 is indeed expressed in normal epithelial cells. We also analyzed skin samples from two younger (under 40) and two older (over 80) non-melanoma skin cancer patients and found that epithelial LINE-1 expression was much higher in malignant cells and somewhat higher in the older patients. Finally, we looked at hippocampal cells from two young (4 month old) and two old (24 month old) mice, and found LINE-1 to be expressed in mouse hippocampal neurons, with modestly increased expression in the older neurons.

## Results

### 3’ targeted scRNA-seq is not appropriate for LINE-1 quantitation

We tested three scRNA-seq methods for their ability to capture LINE-1 expression: the standard 3’ targeted 10x Genomics method (figure 1A), the standard 10x Genomics 5’ method (figure S2) and a 5’ library prep followed by 150 base pair (bp) paired end sequencing (figure 1B). LINE-1 was quantified through a simple method based on our MapRRCon^34^ pipeline (Figure 1C). Reads are fed into a custom Cell Ranger index that includes a masked hg38 genome with all L1HS and L1PA sequences removed and replaced by consensus sequences available from Dfam. Reads aligned to the L1HS consensus are filtered for alignment quality and then unique molecular identifiers (UMIs) are counted.

**Figure 1:**
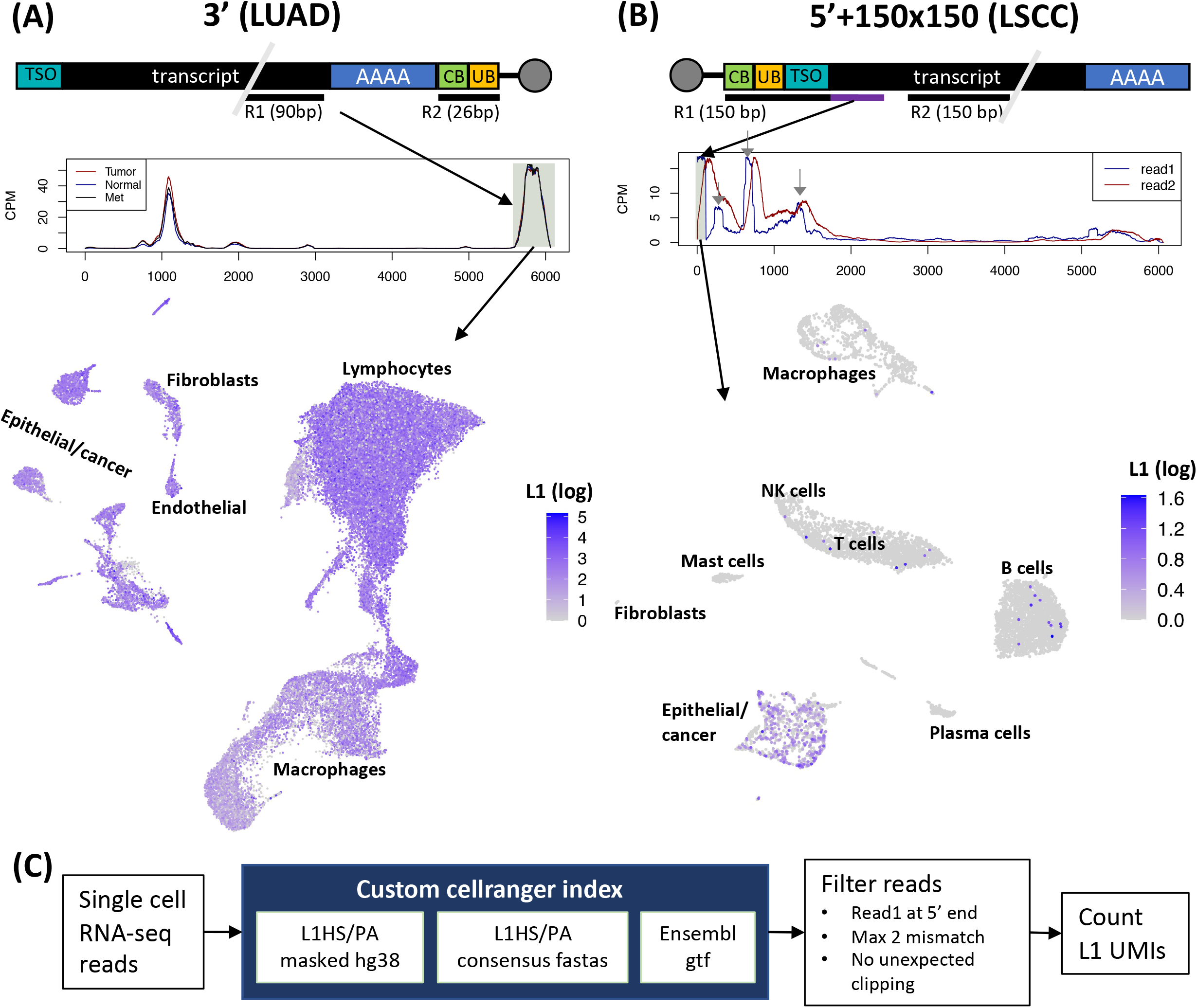
5’ but not 3’ single cell RNA-seq can capture LINE-1 expression. (A) 3’ targeted single cell RNA-seq fails to capture LINE-1 expression in a lung adenocarcinoma (LUAD) tumor. Top: Schematic of the 3’ 10x Genomics sequencing strategy. mRNAs are captured at the 3’ end by bead bound poly dT primers attached to a cell specific barcode (CB, 16bp) and a molecule specific barcode (UB, 10bp). The cDNA is fragmented and a read pair is sequenced with a 90bp read 1 (R1) falling at the 5’ end of 3’ fragment and a 26 bp read 2 (R2) covering only the barcodes. Middle: Coverage of reads aligned to the LINE-1 consensus. A lack of difference between tumor/met and normal immediately suggests that this data fails to accurately capture LINE-1 expression. Bottom: UMAP embedding of all cells colored by LINE-1 “expression.” The presence of large numbers of LINE-1 aligning reads in each cell type is further evidence that 3’ targeted single cell RNA-seq is ineffective for LINE-1. (B) 5’ targeted single cell RNA-seq with 150 bp paired end reads successfully captures LINE-1 expression in lung squamous cell carcinoma (LSCC). Top: Schematic of the 5’ 10x Genomics sequencing strategy (with 150 bp paired reads). mRNAs are captured at the 5’ end by a bead bound template switch oligo (TSO) attached to a cell specific barcode (CB, 16bp) and a molecule specific barcode (UB, 10bp). The cDNA is fragmented and a read pair is sequenced with a 150 bp read 1 (R1) that covers the bar codes (26bp), the TSO and, *crucially*, the exact 5’ end of the transcript sequence (purple section) and a 150 bp read 2 that falls at the 3’ end of the 5’ fragment. Middle: Coverage of reads aligned to the LINE-1 consensus. Transcripts from the LINE-1 promoter is represented by the read 1 peak at the 5’ end. Bottom: UMAP embedding of all cells colored by LINE-1 expression from the 5’ read 1 peak. The known LINE-1 expression pattern – specific to cancer cells – is recapitulated. (C) 5’ sc-L1 pipeline. Single cell rna-seq reads are aligned to custom Cell Ranger index. Reads are filtered for alignment quality and a read 1 starting within 20bp of the 5’ end of L1Hs. LINE-1 unique molecular identifiers (UMIs) are identified from the unique barcodes (UB) and assigned to cell based on the cell barcode (CB).

LINE-1 expression (i.e. RNA transcript accumulation driven by the internal LINE-1 5’ promoter), was vastly overestimated when we reanalyzed normal lung, tumor, and metastases (met) samples from lung adenocarcinoma (LUAD) patients sequenced by the 3’ method^35^. In this data, only the 90bp read 1 carries any information about the captured RNA molecules (figure 1A, top). Read 1 sequences aligned to the L1HS consensus in two peaks (figure 1A, middle). The smaller peak likely reflects a cryptic polyA site in the LINE-1 sequence (an AAUAA motif appears at the peak’s 3’ end.) The second, larger peak occurred as expected at the 3’ end of LINE-1. However, there was no difference between normal, tumor and met, despite considerable evidence showing that LINE-1 is upregulated in cancer cells^36^.

We then mapped reads from this second peak into a UMAP embedding and found high LINE-1 “expression” in nearly every cell type (figure 1A, bottom), further indicating that this method does not reflect the known pattern of tumor LINE-1 expression. LINE-1 is present in the human genome at a very high copy number, but most of these copies are inactivated by mutation and/or 5’ truncation. However, LINE-1s, including the half million or so copies that lack promoter activity, are frequently transcribed into non-LINE-1 transcripts, and these non-LINE-1 specific “passive” transcripts often vastly exceed “active” LINE-1 expression^37,38^. Any of these passive transcripts that are poly-adenylated at LINE-1’s 3’ polyadenylation site will be indistinguishable from active LINE-1 expression in 3’ targeted datasets. Therefore, it is not entirely unexpected that a 3’ targeted sequencing method is not effective at measuring LINE-1 expression.

### 5’ targeted 10x Genomics sc-RNA (with an extended read 1 length) captures LINE-1 expression

Because passive LINE-1 transcripts initiate upstream of the element, they can be filtered out by using a 5’ targeted method. 10x Genomics methods use a transcript switch oligo (TSO) to identify RNA molecules that have been fully reverse transcribed into cDNA. Polymerase reaching the 5’ end of the RNA molecule will change templates at the TSO, yielding a molecule where the TSO sequence is affixed to the 5’ end of the fully reverse transcribed cDNA (figure 1B, top). Thus, if we perform sequencing that is 100 or 150 bp paired end, we will sequence the TSO/cDNA junction, allowing us to precisely identify LINE-1 RNAs that start within a few bases of the 5’ end (i.e. result from LINE-1 promoter activity; see purple segment in figure 1B top). To test this assertion, we aligned reads from a 10x genomics example dataset that includes 150 bp paired end reads from a different – lung squamous cell carcinoma (LSCC) – tumor^39^. This yielded a large peak of read 1 sequences mapping within 20 bases of the 5’ end of LINE-1 (shaded peak in figure 1B, middle) as well as a few peaks that are likely not representative of LINE-1 promoter activity (gray arrows in figure 1B, middle). When we only count UMIs from read pairs that have a read 1 in this peak – a method we call 5’ scL1seq – LINE-1 expression is almost entirely restricted to the epithelial/cancer cluster, reflecting abundant evidence that LINE-1 is upregulated in cancer cells^14,21,36^.

The sequencing 2×150bp template described above differs from the default template that is currently employed for 10x genomics sequencing (5’ v2 chemistry). In the default template, read 1 is 26bp and only covers the cell and UMI barcodes, while read 2 is 90 bp and provides information about which transcript has been captured (figure S2, top). Thus, only the read 2 coverage (red in figure 1B, top) is recovered. However, while the read 1 coverage shows two peaks in the first 500bp (shaded peak and first gray arrow in figure 1B, middle), the read 2 peaks corresponding to the proper LINE-1 expression peak and to the second peak blend together, making it impossible to fully disentangle read 2 sequences that come from proper LINE-1 transcripts. We tested the standard 10x genomics 5’ single cell RNA-seq template on HeLa cells +/-an L1RP plasmid. In the L1RP-cells, we observed a peak of reads near the 5’ end of LINE-1 that cannot be filtered out without losing sensitivity in the L1RP+ cells (figure S2). Thus, it is important to sequence a 100+bp read 1 that will cover the precise TSO/transcript junction.

### LINE-1 is expressed in normal human epithelial cells

Having developed a high throughput method to measure LINE-1 RNA expression in single cells, we returned to the primary question motivating this study: how much LINE-1 expression occurs in normal, adult somatic cells? To that end, we reanalyzed 5’ targeted single cell RNA-seq data sequenced using 150 bp paired end reads from 15 healthy human tissues: bile duct, bladder, blood, esophagus, heart, liver, lymph node, bone marrow, muscle, rectum, skin, small intestine, spleen, stomach and trachea^40^. Fibroblasts, endothelial, muscle, and immune cells clustered by cell type regardless of tissue origin and showed almost no LINE-1 expression. However, epithelial cells, which clustered by tissue of origin, showed clear evidence for LINE-1 expression (figures 2A, S2). The presence of LINE-1 expression in normal epithelial cells is reminiscent of the fact that frequent retrotransposition events have been identified in epithelial cell derived tumors (carcinomas), but retrotransposition is infrequent or absent from cancers derived from blood and brain^21,41,42^.

**Figure 2.**
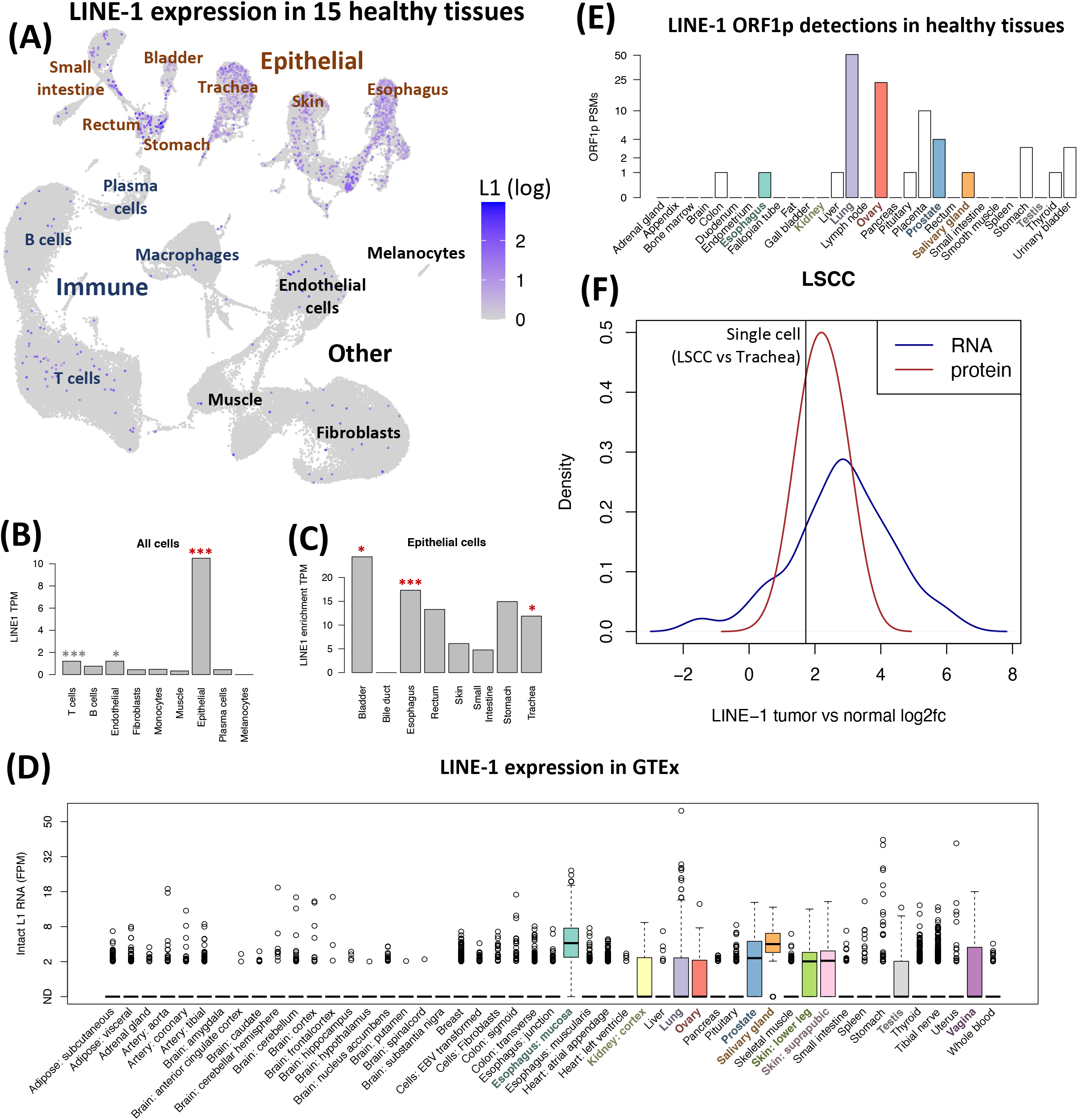
LINE-1 is expressed in normal epithelial cells. (A) UMAP of LINE-1 RNA expression in single cells from 15 healthy tissues of a single healthy donor. Epithelial cells are labeled by tissue of origin. (B) LINE-1 expression by cell type. ***: p<0.001 for enrichment in this cell type, *:0.01<p<0.05. Red indicates comparison to all other cells, gray indicates comparison to other non-epithelial cells. (C) Epithelial cell LINE-1 expression by tissue of origin. ***: p<0.001 for enrichment in this tissue, *:0.01<p<0.05. (D) LINE-1 RNA expression (estimated by L1EM) in bulk tissue samples from GTEx. (E) LINE-1 ORF1p detection in healthy tissues. (F) Comparison of LINE-1 expression in LSCC vs tracheal epithelial cells to bulk tumor/normal comparisons from bulk LSCC samples. Blue and red LINE-1s show the distribution of tumor sample LINE-1 enrichment in RNA and protein respectively. Black LINE-1 indicates enrichment of LINE-1 RNA expression in LSCC epithelial cells compared to tracheal epithelial cells.

To quantify the LINE-1 enrichment in epithelial cells, we fit a negative binomial generalized linear model (nb glm) to the number of LINE-1 UMIs in each cell, using the total number of UMIs per cell and whether a cell is epithelial as predictors and found that LINE-1 expression is ~17x higher in epithelial cells compared to others in this dataset (p<10^-16^, figure 2B red stars). We then fit similar models to non-epithelial cells, separately considering each cell type as a predictor and found that LINE-1 expression is about twice as high in T cells compared to other non-epithelial cells (p=7.2×10^-5^) and about 73% higher in endothelial cells (p=0.01, figure 2B gray stars). We also used the nb glm to compare epithelial cells from different individual tissues samples. The esophagus sample was most significantly enriched for LINE-1 expression (p < 10^-16^), but we also identified increased LINE-1 expression in bladder (p=0.01) and trachea (p=0.02).

Because younger LINE-1 families including L1PA2, 3 and 4 bear very high sequence similarity to L1Hs, it is possible that some of the signal we see here is not from the youngest (and active) L1Hs family, but from recently active but now extinct LINE-1 families. We therefore used our L1EM tool^37^ to map the filtered reads to specific LINE-1 loci. ~94% of reads were assigned to L1Hs family repeats, suggesting that we are indeed mainly measuring L1Hs family LINE-1 expression. ~30% of reads were assigned to a locus in *AUTS2*. This is the only locus in hg38 that exactly matches the L1Hs consensus across the first 500 bases. Thus it may be that our method of filtering reads based on similarity to the consensus biases this analysis toward that locus. Nevertheless, 2 of the 6 loci that were assigned at least 5% of reads (one at 22q12.1 and one at 1p12) are known to generate frequent 3’ transductions in cancer^21^, suggesting that at least some of this normal epithelial cell LINE-1 expression comes from retrotransposon competent loci.

We then wanted to know whether the normal epithelial cell LINE-1 expression that we found in single cells was also detectable in bulk samples. We first applied our L1EM algorithm, which uses the expectation maximization algorithm to separate active LINE-1 expression from passive cotranscription,^37^ to healthy tissue data from GTEx and found that LINE-1 was most often detected in tissues with a significant epithelial cell component. We detected LINE-1 expression (at least one L1Hs locus with > 2 fragments per million / FPM) in at least 25% percent of samples from: esophageal mucosa, kidney, lung, ovary, salivary gland, skin, testis and vagina. On the other hand, LINE-1 expression was rarely detected in tissues lacking in epithelial cells such as adipose (fat), brain, muscle and blood (figure 2B).

We then searched a dataset of particularly deep mass spectrometry proteomics from 29 healthy tissues for spectra matching peptides in the LINE-1 ORF1 protein^43^. At least 4 peptide spectral matches (PSMs) were identified in lung (51), ovary (23), placenta (10) and prostate (4), but ORF1p was not detected in tissues lacking epithelial cells, including bone marrow, brain, fat, lymph node and smooth muscle (figure 2C). This suggests that LINE-1 ORF1p is present in these tissues, but typically lies near or below the detection limit for mass spectrometry proteomics. While LINE-1 RNA expression is present in both tumor and normal epithelial cells, it is still about 3.3x higher in the LSCC epithelial cells, analyzed above, than in the normal trachea epithelial cells. This enrichment is consistent with tumor/normal comparisons of LINE-1 RNA and protein expression in bulk LSCC samples from CPTAC^44^ (figure 2D).

### LINE-1 expression in malignant vs normal cells in younger and older individuals

Because we found LINE-1 expression to be highest in epithelial cells and other studies have found LINE-1 to be most active in epithelial derived tumors, we wanted to perform a direct comparison of LINE-1 expression in matched tumor / normal samples. To that end, we performed matched tumor/adjacent normal 5’ single cell RNA-seq with 100bp paired end reads from four patients: a younger (35 year old) squamous cell carcinoma (SCC) patient, an older (88 year old) SCC patient, a younger (34 year old) basal cell carcinoma (BCC) patient, and an older (84 year old) BCC patient. Consistent with the previous datasets, keratinocytes (keratin producing epithelial cells of the epidermis and hair follicle – KCs) from both the tumor (up 84x over non-KCs, p < 10^-16^) and the normal samples (up 7.6x over non-KCs, p < 10^-16^) most frequently expressed LINE-1 (figure 3A,B). Excluding KCs, we also saw an enrichment in T cells (up 3.6x over other non-KCs, p < 10^-16^) and endothelial cells (up 2.3x over other non-KCs, p < 10^-16^).

**Figure 3.**
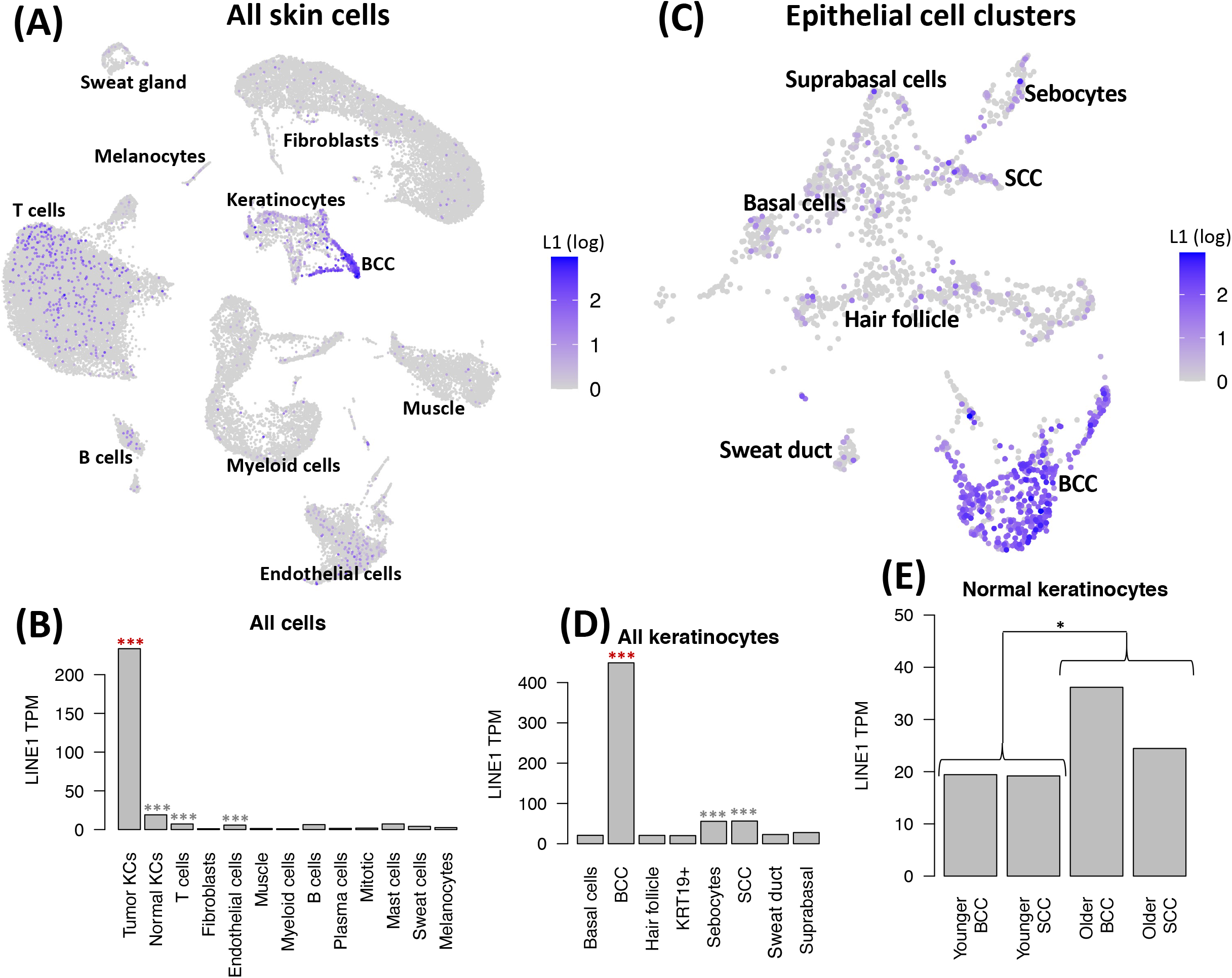
LINE-1 expression in tumor and matched skin from younger and older non-melanoma skin cancer patients. (A) UMAP embedding of all sequenced cells colored by log normalized LINE-1 expression. (B) LINE-1 expression by cell type. ***: p<0.001 for enrichment in this cell type, *:0.01<p<0.05. Red indicates comparison to all other cells, gray indicates comparison to other non-tumor KCs. (C) UMAP embedding of reclustered KCs, colored by log normalized LINE-1 expression. (D) KC LINE-1 expression by subtype. ***: p<0.001 for enrichment in this cell type, *:0.01<p<0.05. Red indicates comparison to all other cells, gray indicates comparison to other non-BCC KCs. (E) Comparison of normal sample KC LINE-1 expression in the under 40 (younger) and over 80 (older) samples.

To get greater detail on LINE-1 expression in KCs, we reclustered the epithelial cells separately (figure 3C). LINE-1 expression was by far the highest in malignant BCC cells (up 15x over other KCs, p < 10^-16^), but was also elevated in SCC cells (up 2.3x over other non-malignant KCs, p=4.6×10^-6^). Unexpectedly, we also found LINE-1 expression to be elevated in sebocytes (up 2.2x over non-BCC KCs, p=8.8×10^-7^, figure 3D).

LINE-1 has been hypothesized to be de-repressed as a result of heterochromatin loss during aging. We therefore wanted to know whether the epithelial cell LINE-1 expression identified here was affected by age. We then used an nb glm model to test a binary age variable (>80 years vs <40 years) as a predictor of LINE-1 expression in our data. We found that advanced age predicted a modest 35% increase in normal KC LINE-1 expression (p = 0.04, figure 3D). However, with only two patients, this analysis lacked the power to fully account for differences in sample cell type make-up. In particular, the older SCC patient included many hair follicle associated KCs that were absent from other samples (figure S1 bottom right).

### LINE-1 is expressed in mouse hippocampal neurons

While we did not see clear evidence for LINE-1 expression in the bulk human brain tissue samples we analyzed above, a number of studies have suggested a role for LINE-1 in normal and diseased brains. We therefore wanted to know whether LINE-1 expression in the brain can be detected at the single cell/nucleus level. While we were not able to find appropriate data from human brain cells, we did identify 150 bp paired-end read 5’ targeted single nucleus RNA-seq data from the hippocampi of two young (4 month old) and two old (24 month old) mice. Because mice have three active LINE-1 families (Tf, Gf and A), we aligned reads to the L1MdTf_I, L1MdGf_I and L1MdA_I consensus sequences^45^. After filtering low quality alignments, we found L1MdTf_I to be most promising for LINE-1 expression (figure 4A): A read 1 peak occurs in the tandem repeat region (at the position where transcription is expected to initiate) and is followed by a broader read 2 peak. A large number of reads that align near the 3’ end of the sequence are likely a form of background and were filtered out from further analyses. Few reads were found to align to L1MdA_I and read alignments to L1MdGf_I did not form as clear a peak at the 5’ end. We then counted L1MdTf UMIs and found that LINE-1 expression was highest in neurons, particularly those of the *cornu ammonis* (CA / hippocampus proper, figure 4B). Overall, neuronal LINE-1 expression was about twice glial LINE-1 expression (p=2.2×10^-8^), and neurons of the *cornu ammonis* expressed about 50% more LINE-1 than those of the dentate gyrus (p=4.3×10^-4^, figure 4C). We then used the nb glm model to test whether advanced age was predictive of increased LINE-1 expression in neurons. This yielded a modest effect similar in scale to what we observed in the skin samples: LINE-1 expression was 25% higher in older neurons (p=0.039, figure 4D).

**Figure 4.**
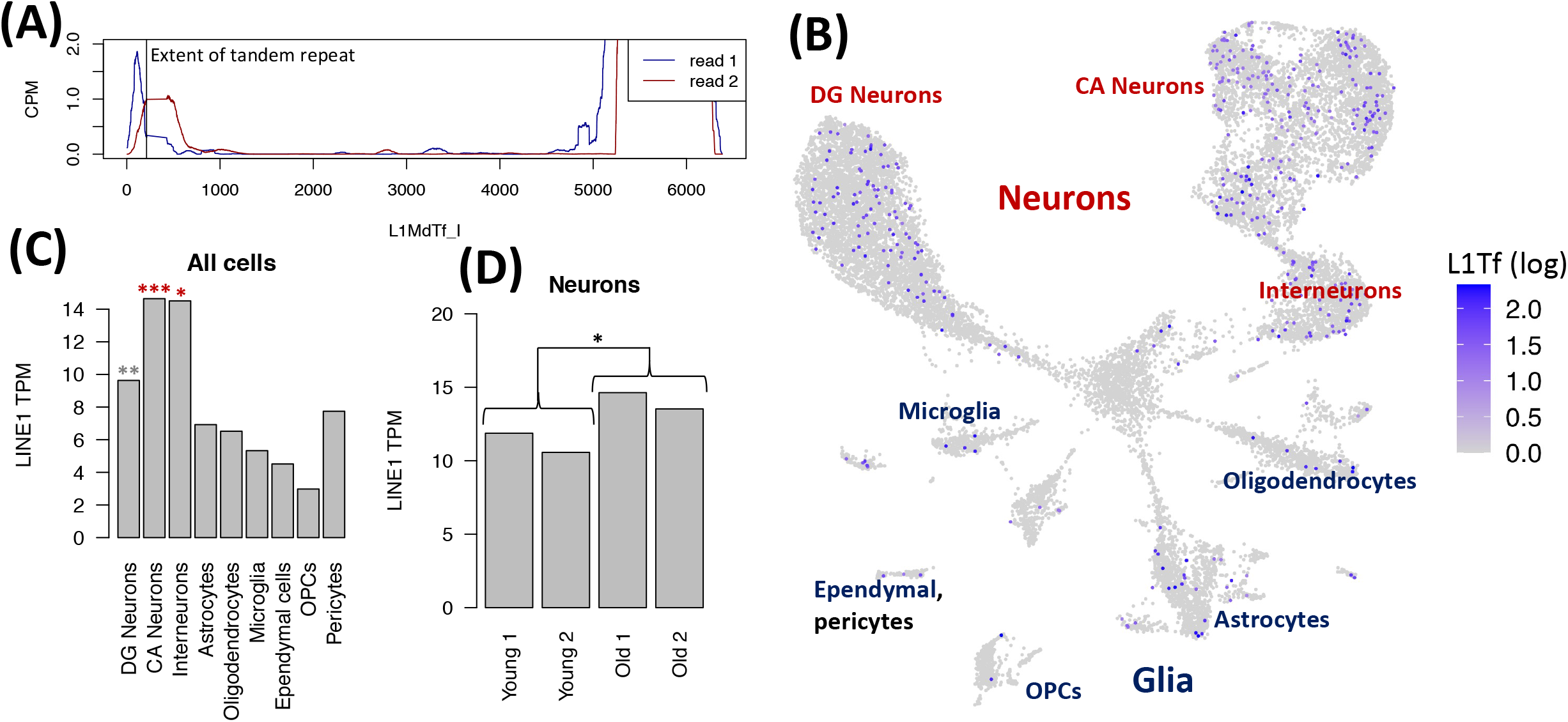
LINE-1 (L1MdTf) expression in the mouse hippocampus by single nucleus RNA-seq. (A) Read coverage across the L1MdTf_I consensus. (B) UMAP embedding of mouse hippocampal cells colored by log normalized L1MdTf expression. (C) LINE-1 expression by cell type. ***: p<0.001 for enrichment in this cell type, ** 0.001<p<0.01, *:0.01<p<0.05. Red indicates comparison to all other cells, gray indicates comparison excludes CA neurons. (E) Comparison of neuronal LINE-1 expression in 4 month old mice (young) to 24 month old mice (old).

## Discussion

While very strong evidence from both RNA and protein^36,46,47^ shows that LINE-1 is highly expressed in many tumors, the extent of its expression in normal cellular contexts is much less well understood. New evidence has linked retrotransposons broadly or LINE-1 specifically to a variety of non-cancer diseases, including age-related inflammation^16,32^, neurodegenerative diseases^48–50^ and autoimmune disorders^51,52^. This makes it all the more important that we have a clear and accurate understanding of where and when LINE-1 expression is driven from its own promoter in human cells. To that end, we developed 5’ scL1seq, a method to quantify LNE-1 expression in single cell RNA-seq data generated on the 10x Genomics platform, and found that the 5’ targeted method combined with 100+bp paired end reads is particularly effective for accurately identifying LINE-1 expression.

When we analyzed data generated by this method for LINE-1, we were surprised to find clear evidence for LINE-1 expression in normal epithelial cells in multiple tissue types. This reflects the fact that LINE-1 activity in cancer is largely limited to tumors of epithelial origin^18–23,41,42^. This relationship between normal and tumor cell LINE-1 expression suggests that LINE-1 de-repression in cancer may not so much be a switch from “repressed” to “expressed” as much as the widening of a pre-existing window of LINE-1 expression. Somatic LINE-1 expression in primary cells prior to tumor development may also have important implications for how cancer develops – particularly in the acquisition of p53 mutations^14^. It is, however, important to note that the quantifications here are only of LINE-1 RNA. Because there are many post-transcriptional mechanisms for LINE-1 regulation^53^, future research is needed to show whether the expression we observe here leads to the translation of LINE-1 proteins and to new retrotransposition events.

We then looked at direct tumor/normal comparisons from non-melanoma skin cancer patients to see how normal LINE-1 expression relates to tumor LINE-1 expression. Despite seeing similar levels of normal LINE-1 expression in basal and suprabasal cells – the normal equivalent for basal (BCC) and squamous cell carcinomas (SCC) respectively – we saw much higher LINE-1 expression in the cancer cells from BCC than SCC. This suggests that LINE-1 expression in tumors is not simply a reflection of the background expression in relevant cell types of origin, but also involves tumor-specific factors, yet to be elucidated.

Because aging is associated with a loss of heterochromatin that can lead to expression of retrotransposons, including LINE-1^16,32,33,54,55^, particularly in senescent cells^16^, we wanted to know whether the epithelial LINE-1 we observe here is an age-related phenomenon. While we did find that tumor adjacent normal skin from two patients over 80 had higher LINE-1 expression than tumor adjacent normal skin from two patients under 40, the effect was modest. Thus it may be that the modest increase with age is due to the increased prevalence of senescent cells, but that there is also another source of epithelial cell LINE-1 expression that is present in both young and old tissues.

Finally, we analyzed nuclei from the mouse hippocampus for LINE-1 expression, where we found it to be expressed in neurons, especially those of *cornu ammonis* (hippocampus proper.) In cancer cell lines, LINE-1 RNPs are primarily cytoplasmic^7^, so it’s possible that the single *nucleus* RNA-seq used to evaluate neuronal expression is measuring a distinct population of LINE-1 mRNAs from those present in malignant and normal epithelial cells.

In this study, we developed a method to accurately quantify LINE-1 expression in tens of thousands of single cells and were able to make significant progress toward answering the question of what constitutes normal LINE-1 expression and how it compares to tumor LINE-1 expression. However, our results also raise the question as to why LINE-1 is more highly expressed in epithelial cells and neurons compared to other cells. These questions highlight how much we still have to learn about how LINE-1 expression is regulated in vivo.

## Methods

### Construction of the cell ranger index

To build the custom reference genome, L1Hs and L1PAx family repeats were masked in hg38 using bedtools maskfasta. L1Hs and L1PAx annotations were downloaded in bed format from the UCSC genome table browser (https://genome.ucsc.edu/cgi-bin/hgTables). Then this masked reference was concatenated with the human L1Hs consensus sequence from repbase^56^ and all available L1PA consensus sequences from Dfam (https://www.Dfam.org/browse?name_accession=L1PA&clade_descendants=true). For transcript annotation, we used the RefSeq GRCh38 annotation in gtf format. A Cell Ranger index was built then with Cell Ranger mkref.

The mouse version of L1-sc was generated in the same manner as human. We first used bedtools maskfasta to mask L1Md sequences (downloaded from the UCSC genome table browser in bed format) from the mm39 mouse genome. We then added L1Md consensus sequences in Dfam (https://Dfam.orgIbrowse?name_accession=L1Md&classification=root;Interspersed_Repeat;Transposable_Element;Class_I_Retrotransposition;LINE;Group-II;Group-1;L1-like;L1-group;L1&clade=10088&clade_descendants=true) as decoy chromosomes. L1MdTf_II/II, L1MdGf_II, and L1MdA_II/II were excluded as they are highly similar to the L1MdTf_I, L1MdGf_I and L1MdA_I consensuses.

### Quantification of LINE-1 UMIs in humans

LINE-1 mRNA UMIs were identified in the Cell Ranger bam file as follows: For the 3’ targeted data, unclipped reads aligning in the final 1kb (but in not the poly A stretch at the 3’ end) of the L1Hs consensus with two or fewer mismatches and no soft clipped reads were checked for the presence of the cell barcode (CB) and UMI barcode (UB) flags. If both were present the UB was added to a python “set” object within a python dictionary with the CB being the key. Once the whole bam file has been read, the number of L1Hs UBs associated with each CB is reported. For the 5’ targeted data from HeLa cells, quantification was performed in the same manner except reads were instead collected from either the first 150 or first 500 bases of the L1Hs consensus. For the 5’ data with 100+bp paired end reads, UMIs were only counted if they have a proper read 1 / read 2 alignment that meets the following criteria: For read 1, the first aligned base must in the first 20 bases of the L1Hs consensus, with less than 20 bases clipped from 5’ end (nb: this number is dependent on how the reads are trimmed), no clipping at the 3’ end, and no more than 2 mismatches. For read 2, neither end can be clipped and no more than 2 mismatches are allowed. To recreate the behavior of Cell Ranger’s aggr function, when combining samples, reads were downsampled to assure even coverage between samples.

### Quantification of LINE-1 in mouse

The mouse L1 promoter has a somewhat different structure from the human version, making it more challenging to quantify 5’ targeted LINE-1 reads in mice. Rather than using the downstream promoter method that allows human LINE-1 to carry its promoter to a new location, the mouse 5’ UTR contains multiple copies of its promoter (i.e. a series of tandem repeats of ~212 bp), with additional copies potentially being generated during insertion (presumably by slippage during reverse transcription). Given this repetitive 5’ UTR structure, it was not obvious where the true “Start” site was located in the monomer. We analyzed the monomer sequence from the Orleans Reeler L1MdTf insertion and looked for a match to the initiator consensus sequence Inr^57,58^. We found a single perfect match to Inr, and used this as our landmark as the inferred start site of L1MdTf initiation. When we aligned reads to this modified consensus, we found a read 1 peak at the initiator sequence (figure 4A). However, the exact mouse LINE-1 TSS was not as precise as it was in human LINE-1, so we included all read pairs with a read 1 fall within a tandem repeat in our mouse LINE-1 quantification. The same clipping and mismatch filters were applied: no more than 2 mismatches and no clipping except up 20 bases at 5’ end of read 1.

### Single cell RNA-seq clustering and cell identification

A count matrix was built using Cell Ranger count and the index described above (default parameters). Cell ranger aggr was used to merge related runs. Clustering and UMAP embedding was performed in Seurat according to the PBMC 3k tutorial (https://satijalab.org/seurat/v3.2/pbmc3k_tutorial.html). Cells with more than 25% MT-RNA, or less than 1000 RNA molecules detected were removed. Clusters showing markers for disparate cell types were taken to be doublets and removed. Human cell types were then identified using marker genes: CD3E for T cells, MS4A1 for B cells, MZB1 for plasma cells, GNLY for NK cells, LYZ for macrophages, KIT for mast cells, COL1A2 for fibroblasts, ACTA2 for muscle, PLVAP for endothelial cells, MLANA for melanocytes, and keratin (KRT) genes for epithelial cells (supplemental figures S2–S5). For keratinocyte clusters we used: KRT5 for basal cells, KRTDAP for suprabasal cells, KRT28/19 for hair follicle associated cells, PPARG for sebocytes and KRT77 for sweat duct cells (figure S6). Malignant cells were identified as CNV+ cells by inferCNV figure S7). The following markers were used in the mouse hippocampus: Gfap for astrocytes, Tmem119 for microglia, Myrf for oligodendrocytes, Rbfox3 for neurons, Cspg4 for OPCs, Cldn5 for ependymal cells, and Slc6a13 for vascular cells. Interneurons were distinguished by Slc6a1 expression, DG neurons by C1ql2, and CA neurons by Neurod6 and Dkk3^59^ (figure S8). UMAP plots were made using the DimPlot and FeaturePlot functions in Seurat.

### 5’ scRNA-seq from HeLa +/-L1RP

#### Cell line and plasmid

HeLa-M2 cells (HeLa cells with a reverse tetracycline-controlled transactivator gene coding for rtTA2-M2, a gift from Gerald Schumann, Paul Ehrlich Institute) were cultured in DMEM supplemented with 10% FBS (Gemini #100–106) and 1x Penicillin-Streptomycin-Glutamine (ThermoFisher #10378016). The cells were routinely tested for *Mycoplasma* by PCR detection of conditioned medium. The L1rp reporter construct - pLAK002 (tet-inducible promoter-L1rp-GFP-AI) plasmid was introduced into HeLa-M2 cells by plating 0.2 × 10^6^ cells per well in a 6-well plate the day before transfection. The following day, cells were transfected with pLAK002 using FuGENE-HD Transfection Reagent (Promega #E2311), following the manufacturer’s protocol. 48 h post-transfection, 1 μg/ml of puromycin was added. The following day, cells were trypsinized and replated in a 10 cm plate with 1 μg/ml of puromycin added. Puromycin was replenished every other day for 14 d until no cell death was observed.

#### Retrotransposition assay in 6-well plates

Once selection was complete, cells were trypsinized and plated at 2.0 × 10^5^ cells per well in a 6-well plate. 24 h post cell plating, doxycycline (1 μg/ml) was added to induce L1 expression. After 72 h, the percentage of GFP+ cells was measured by flow cytometry (FACS buffer: PBS + 2% FBS) to confirm L1 retrotransposition and samples were submitted for scRNAseq.

#### Single-cell RNAseq

Cells were detached from the plate with TrypLE^™^ Express Enzyme (ThermoFisher #12604013) following the manufacturer’s protocol and then resuspended in DMEM at 0.015 × 10^6^ cells per 38.7μL and submitted to the NYULH Genome Technology Core Facility. scRNA-seq was performed using the 10x Genomics system using 5’ gene expression library.

### Negative binomial generalized linear models

Because LINE-1 expression is low in normal cells, there are usually only a few LINE-1 UMIs detected in a single cell. As a result, LINE-1 detection is biased toward cells with a greater number of UMIs, causing the popular wilcoxon rank sum test on log normalized expression to, in some cases, yield biased results. We therefore chose to estimate LINE-1 expression using a negative binomial generalized linear model with log link function:

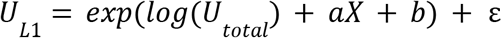

where *U_L1_* is the number of LINE-1 UMIs in a cell, *U_total_* is the total number of UMIs in that cell, *X* is a predictor (such as cell type or age), *a, b* are parameters to be learned, and ε is the negative binomial error term. These models were fit using the glm.nb function in the MASS package for R, using the “offset” function to ensure that the coefficient on *log*(*U_total_*) has a fixed value of 1. P values were estimated by anova. Under this model, transcripts per million (TPM) can be estimated as:

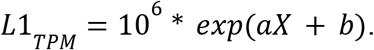

### Single cell suspension derived from patient tumors and matched normal

Patient tumors were processed as described previously^60^. Adjustments are described: tumors were obtained on the day of Mohs micrographic surgery and washed in cold DMEM [Gibco 11995-065] supplemented with 10% FBS [Thermo Scientific SH30910.03]. Any fat was removed from tumor samples using a scalpel. Then, tumors were then cut into small pieces using a razor and resuspended in 10 mL of DMEM/10% FBS, 10 mg/mL collagenase II [Sigma C2674] and 10 U DNase I [Sigma 00453869] for 20 min at 37 °C. After incubation, the suspension was vortexed 1× for 30 s followed by pipetting sequentially through 25, 10, and 5 mL pipettes for 1 min each. Next, the cell suspension was filtered through a 70-μm filter [Fisher 22363548] and spun at 300 g for 5 min. After spinning, the supernatant was removed, and ACK red blood cell lysis was performed according to the manufacturer’s protocol as needed [Gibco a10492-01].

### Single-cell library construction and 5’ sequencing from patient tumors and matched normal

The cellular suspensions were loaded on a 10x Genomics Chromium instrument to generate single-cell gel beads in emulsion (GEMs). Approximately 10–12 x 10^3^ cells were loaded per channel. Single-cell RNA-Seq libraries were prepared using the following single cell 5’ reagent kits: Chromium™ single cell 5’ Library & Gel Bead Kit, PN-1000006 and chromium single cell VDJ enrichment kit for human T cells (PN-100000). Libraries were run on a NovaSeq 6000 SP (SCC) or S1 (BCC) flow cell (depending on the number of samples per run).

### Mouse hippocampus single nucleus RNA sequencing

For snRNA-seq the hippocampus was dissected from the brains of 4-month old and 24 month old C57BL/6 mice. A total of 4 animals were used in each age group. The hippocampi from two mice were pooled together into one sample, and the other two mice were pooled in another sample. Nuclei were isolated from minced hippocampi tissue using the Nuclei PURE Prep Nuclei Isolation Kit with a Dounce B homogenizer. Samples were subjected to a sucrose gradient, and nuclei were further purified and counted. We targeted 5,000 nuclei per sample to load onto a 10x Chromium chip using VD(J) chemistry. We targeted 50,000 sequence reads per nuclei on an Illumina HiSeq device.

## Data and code availability

3’ lung adenocarcinoma data was accessed from the SRA database: PRJNA510251.5’ lung squamous cell data is provided as example data from 10x genomics (https://www.10xgenomics.com/resources/datasets/nsclc-tumor-1-standard-5-0-0). SRA bioproject accession numbers for the other datasets are as follows: PRJNA510251 for 3’ lung adenocarcinoma patient data; PRJNA670909 for 5’ normal tissues data; PRJNA781454 for non-melanoma skin cancer patient data; PRJNA677926 for 5’ mouse hippocampus data. Scripts and readme necessary to run 5’ scL1seq on other datasets can be found on github: https://github.com/wmckerrow/5pL1sc.

## Acknowledgments

We would like to thank Martin S Taylor for his helpful suggestions to improve figure clarity.

---

**Figure S1.**
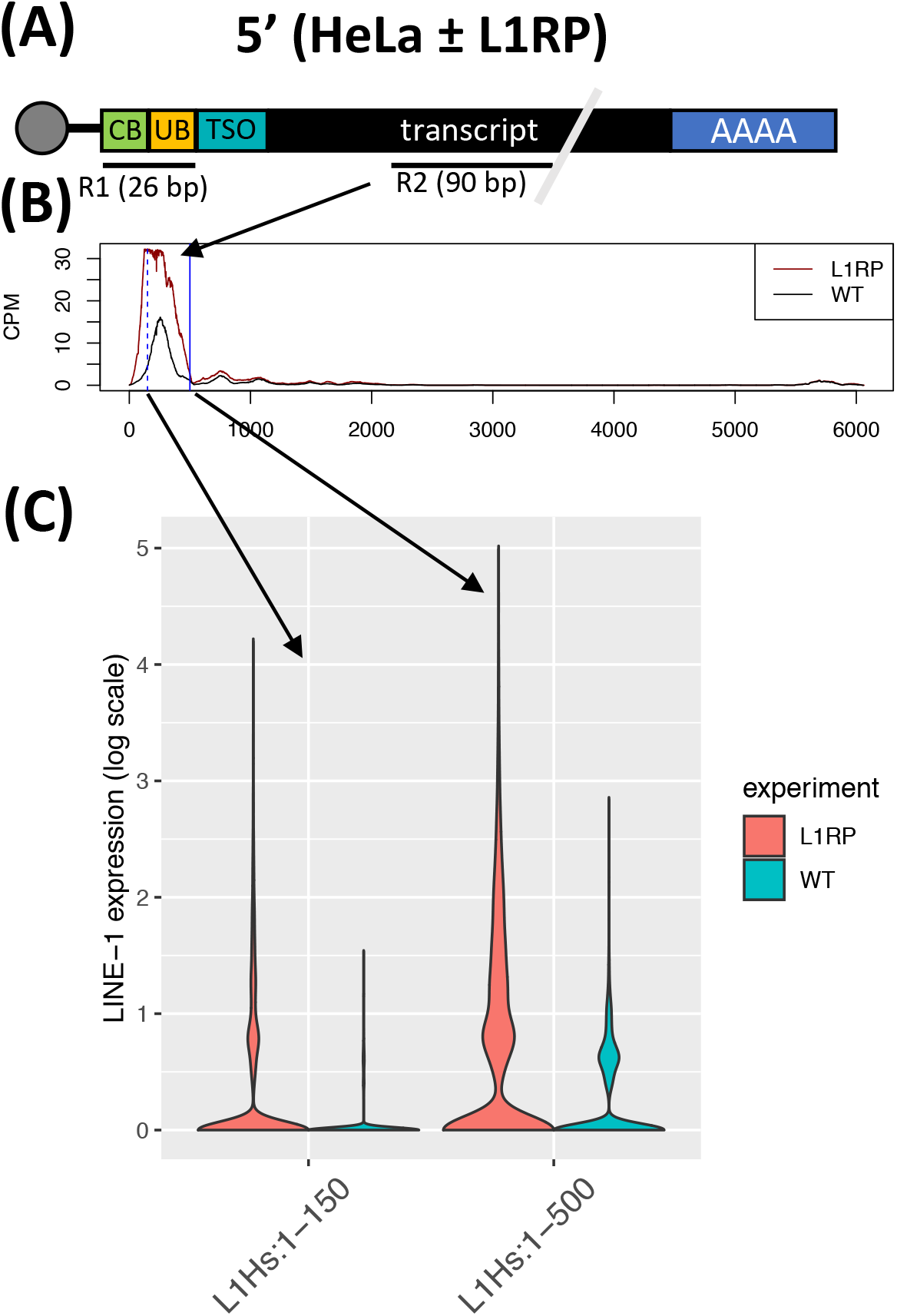
HeLa cells ± L1RP (a LINE-1 overexpression vector) sequenced with the standard (26/90bp) paired end layout. (A) Schematic of sequencing strategy. Exactly as in figure 1A, except that read 1 is only 26 bp and thus only covers the cell (CB) and unique (UB) barcodes and read is only 90 bp long. (B) Coverage of reads along the L1Hs consensus. Collecting reads aligning with the first 150 bp (blue dashed line) will filter out almost all L1Hs aligning read 2s in the L1RP-control. Allowing reads to fall anywhere in the first 500 bp (solid blue LINE-1 will recover all LINE-1 expression in the L1RP+ cells. (C) Violin plots comparing LINE-1 expression in L1RP + vs – HeLa cells using either the 150 or the 500 bp cutoff.

**Figure S2.**
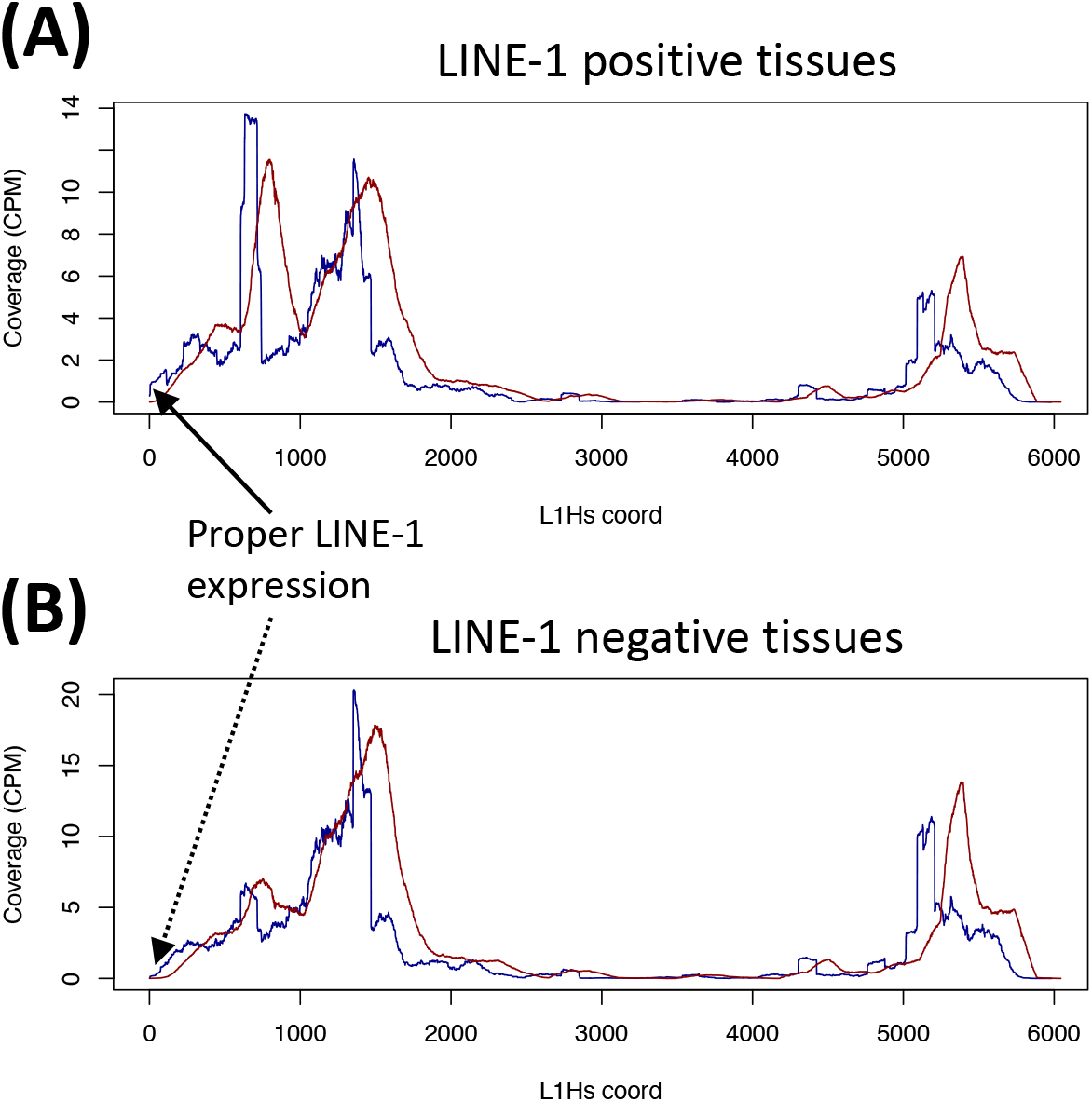
Coverage of L1Hs aligning reads in (A) normal LINE-1+ (bladder, esophagus, rectum, skin, small intestine, stomach, trachea) and (B) LINE-1-tissues (blood, bile duct, heart, liver, lymph node, marrow, muscle, spleen). A small peak of read 1s starting at the 5’ end of LINE-1 can be seen in the LINE-1+, but not the LINE-1-tissues.

**Figure S2:**
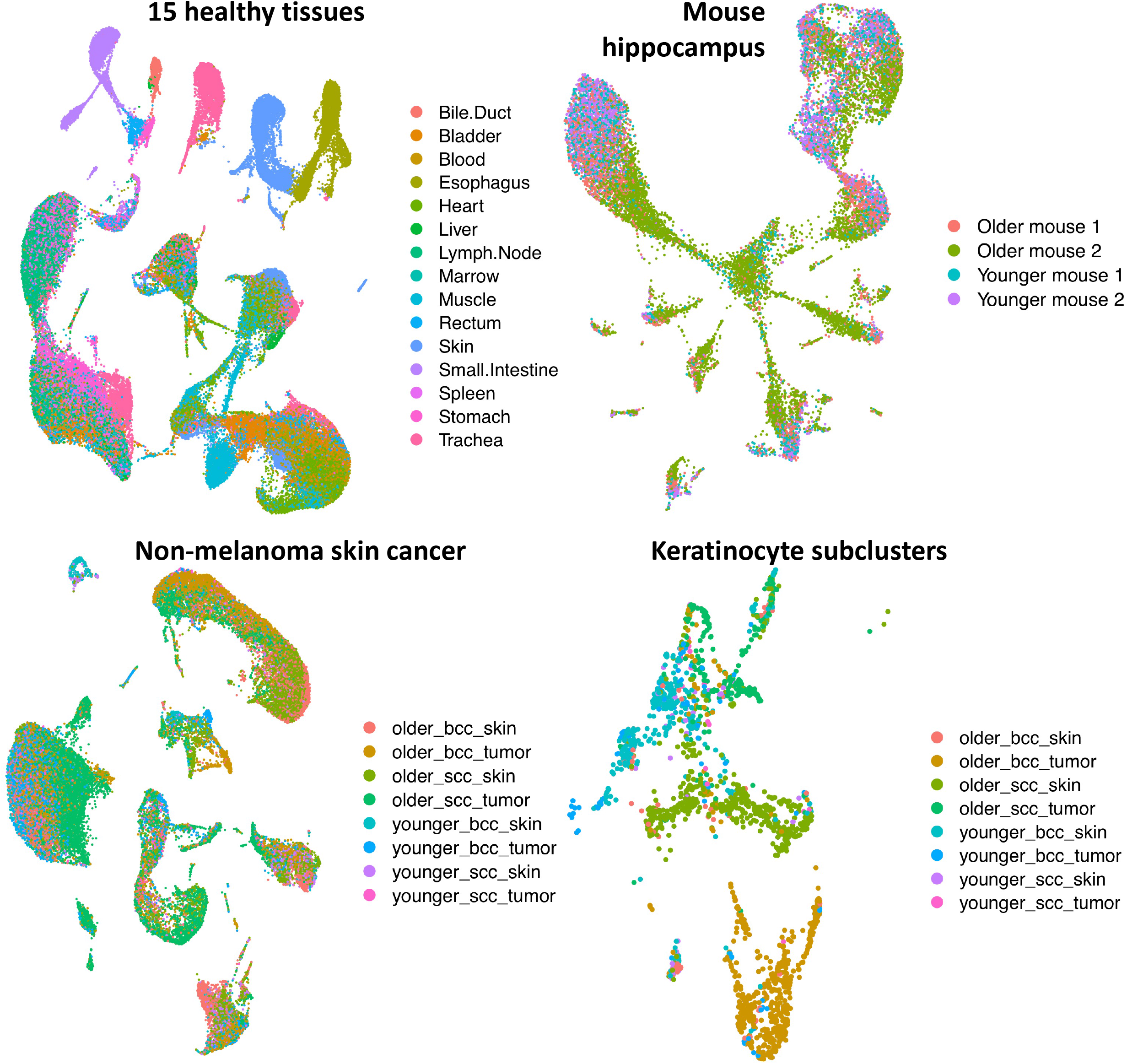
UMAPs colored by sample of origin.

**Figure S2:**
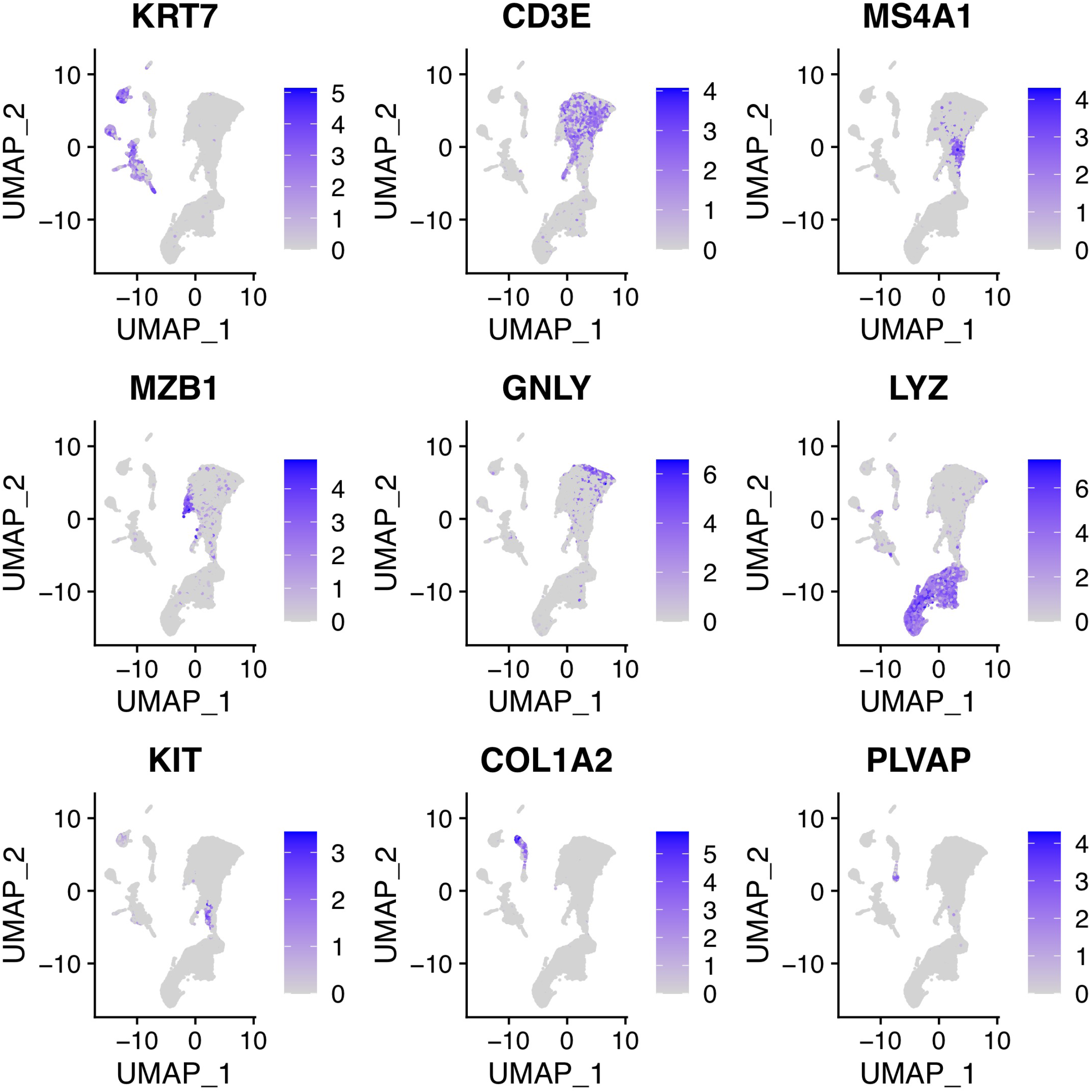
Marker gene expression for lung adenocarcinoma tumor, metastasis and normal cells. KRT7 = epithelial/cancer, CD3E = T cells, MS4A1 = B cells, MZB1 = plasma cells, GNLY = NK cells, LYZ = macrophages, KIT = mast cells, COL1A2 = fibroblasts, PLVAP = endothelial cells.

**Figure S3:**
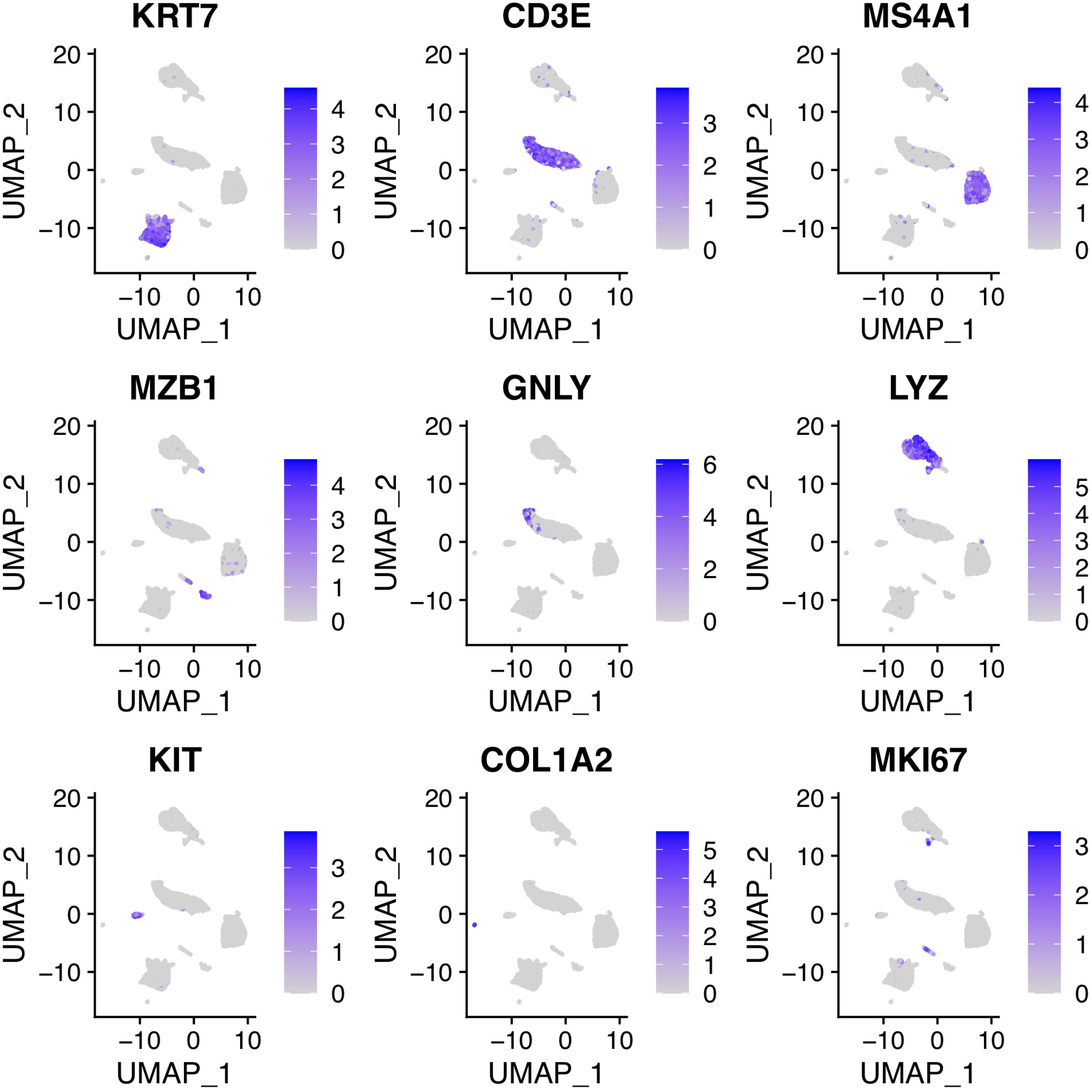
Marker gene expression for lung squamous cell cancer tumor cells. KRT7 = epithelial/cancer, CD3E = T cells, MS4A1 = B cells, MZB1 = plasma cells, GNLY = NK cells, LYZ = macrophages, KIT = mast cells, COL1A2 = fibroblasts, MKI67 = mitotic cells.

**Figure S4:**
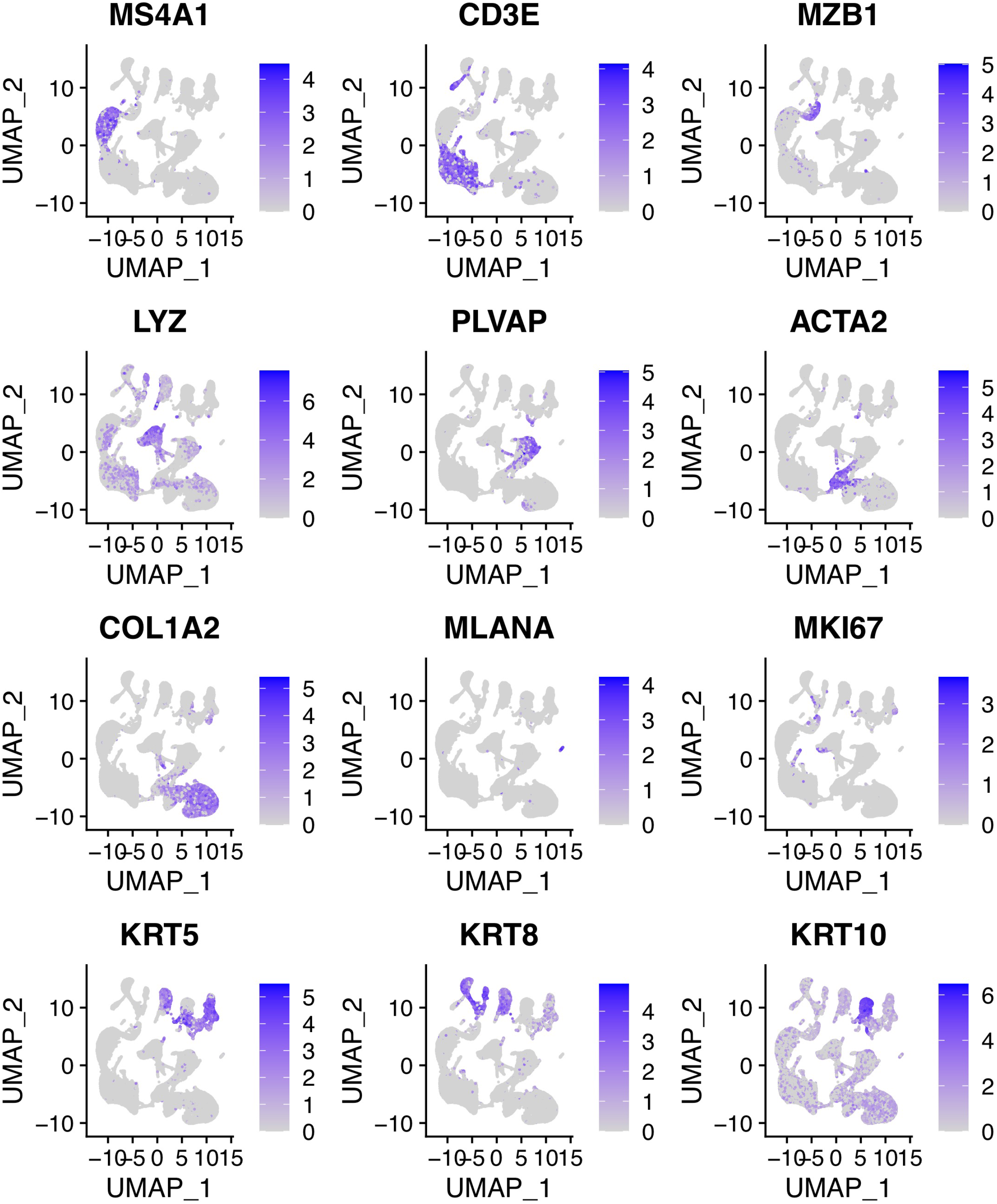
Marker gene expression for normal tissue cells. MS4A1 = B cells, CD3E = T cells, MZB1 = plasma cells, LYZ = macrophages, PLVAP = endothelial, ACTA2 = muscle, COL1A2 = Fibroblasts, MLANA = melanocytes, MKI67 = mitotic, KRTX = epithelial.

**Figure S5:**
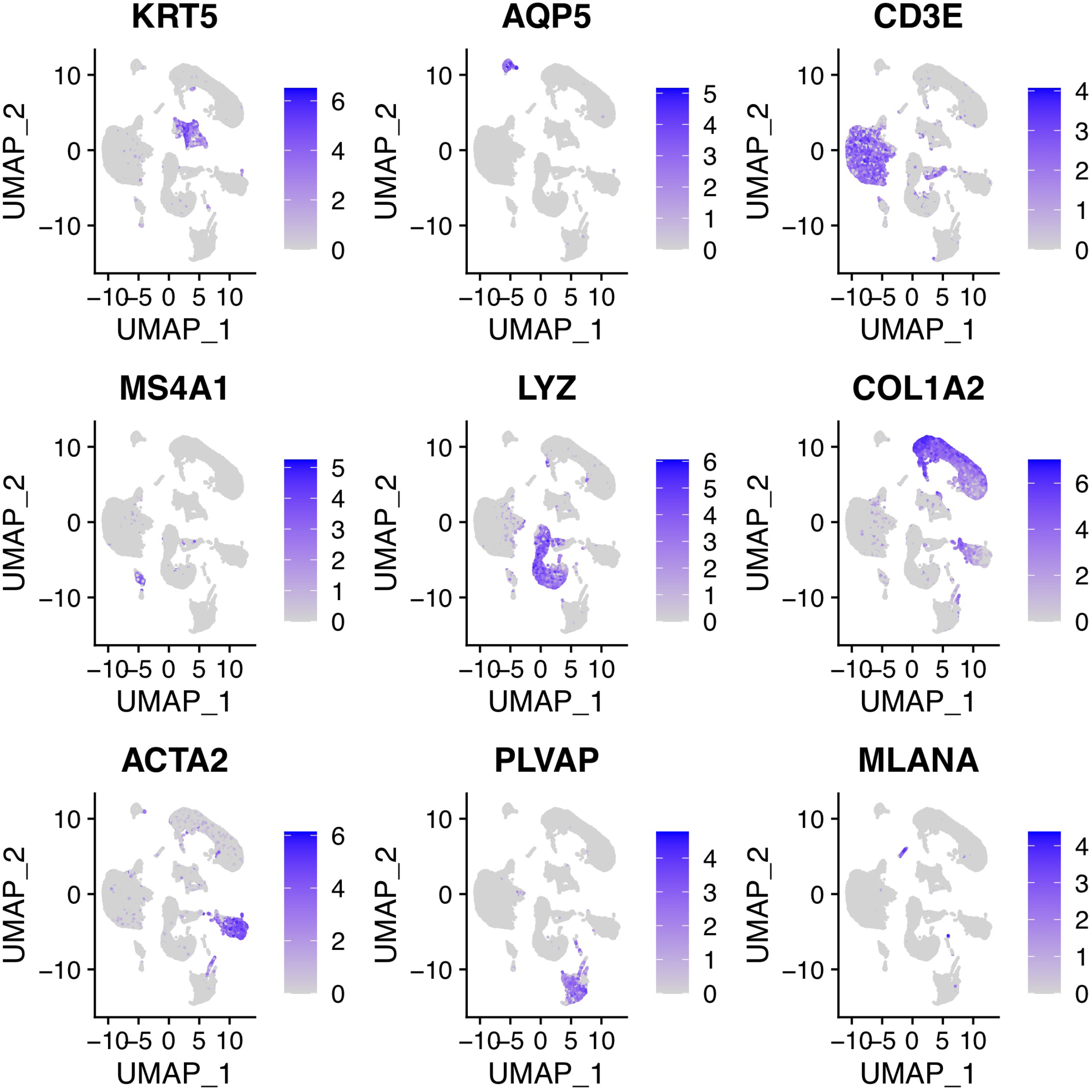
Marker gene expression for non-melanoma skin cancer patients.

**Figure S6:**
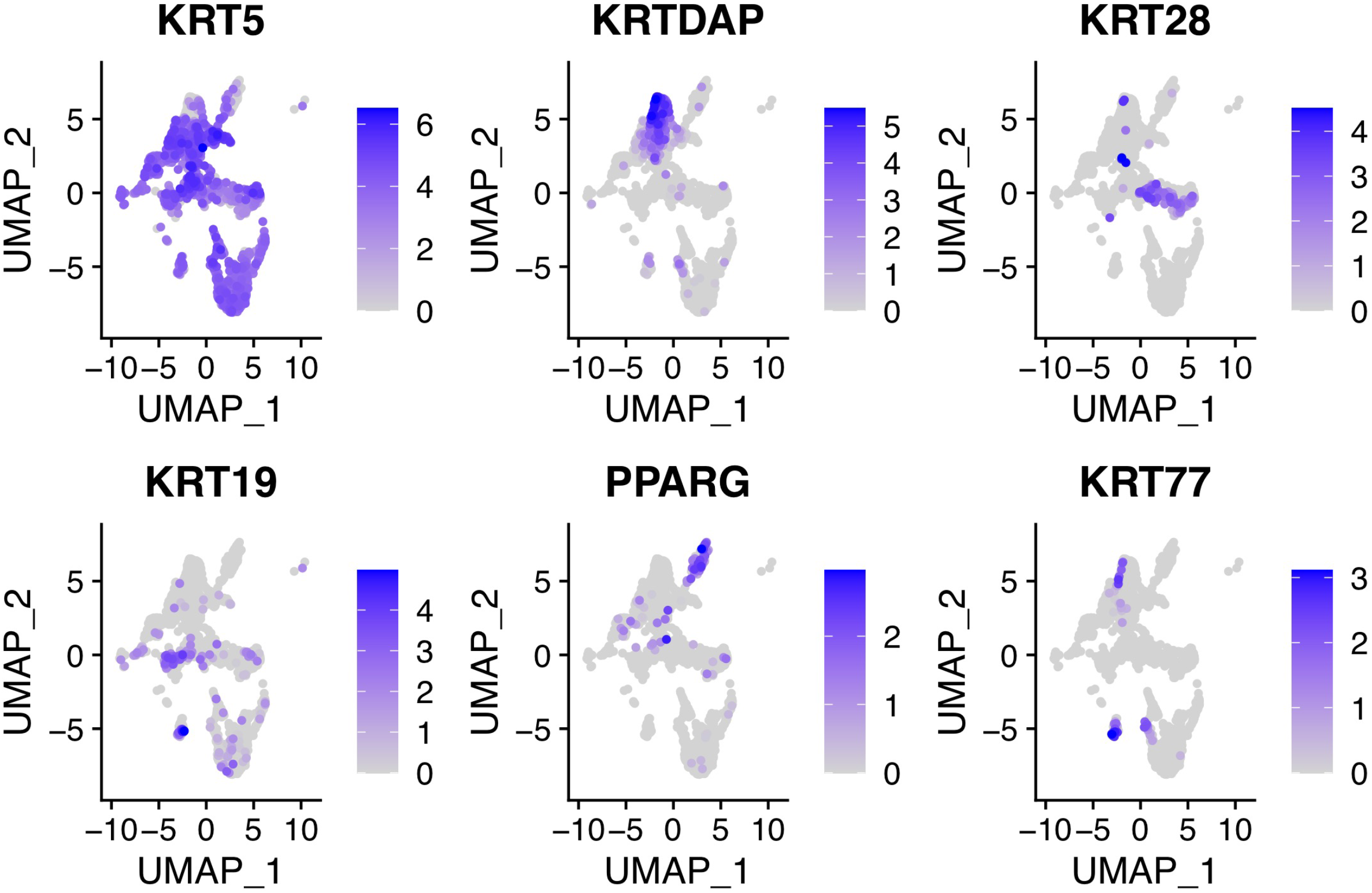
Marker gene expression for non-melanoma skin cancer patients.

**Figure S7:**
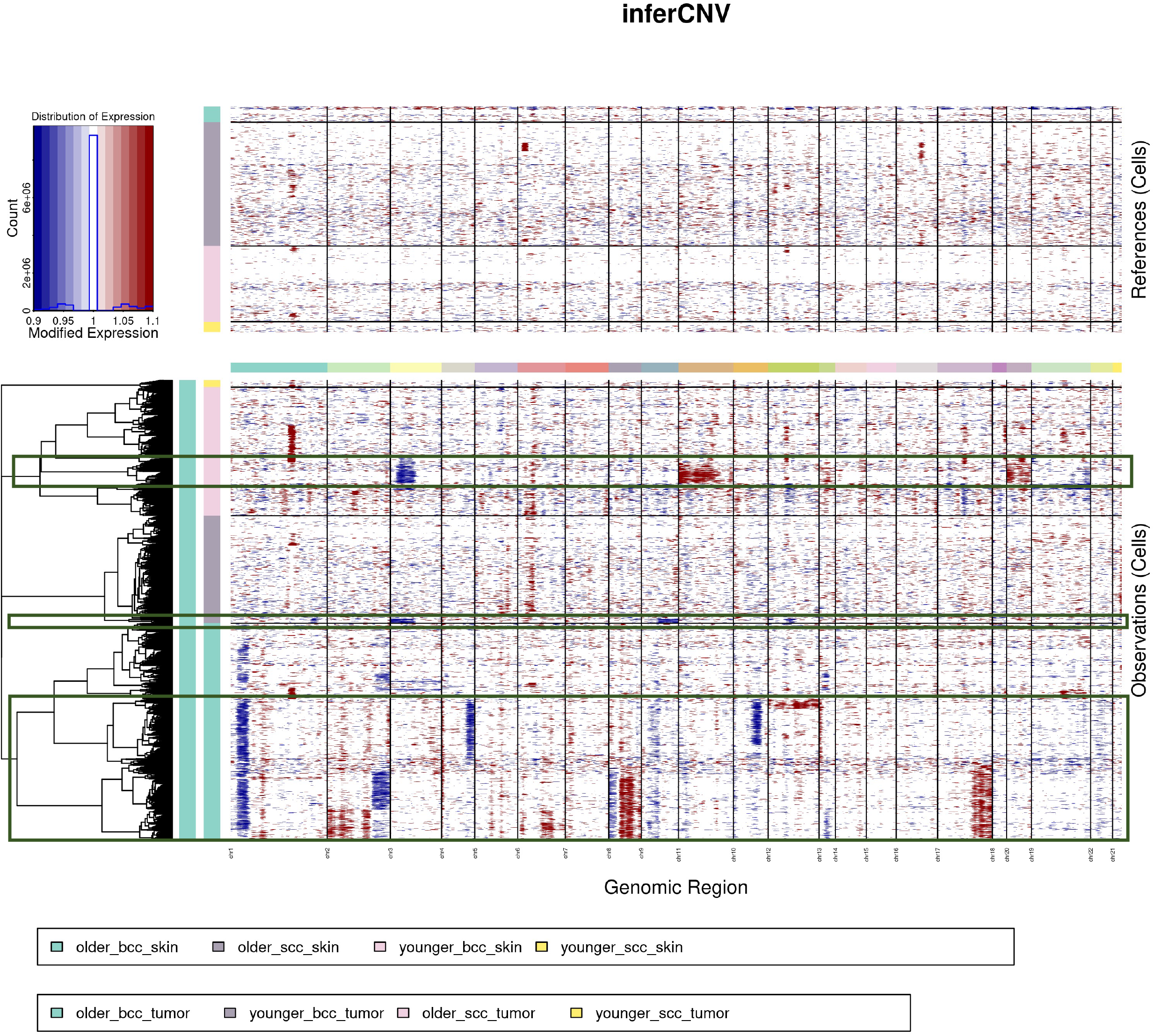
Output of inferCNV. CNV+ cells are highlighted by green boxes.

**Figure S8:**
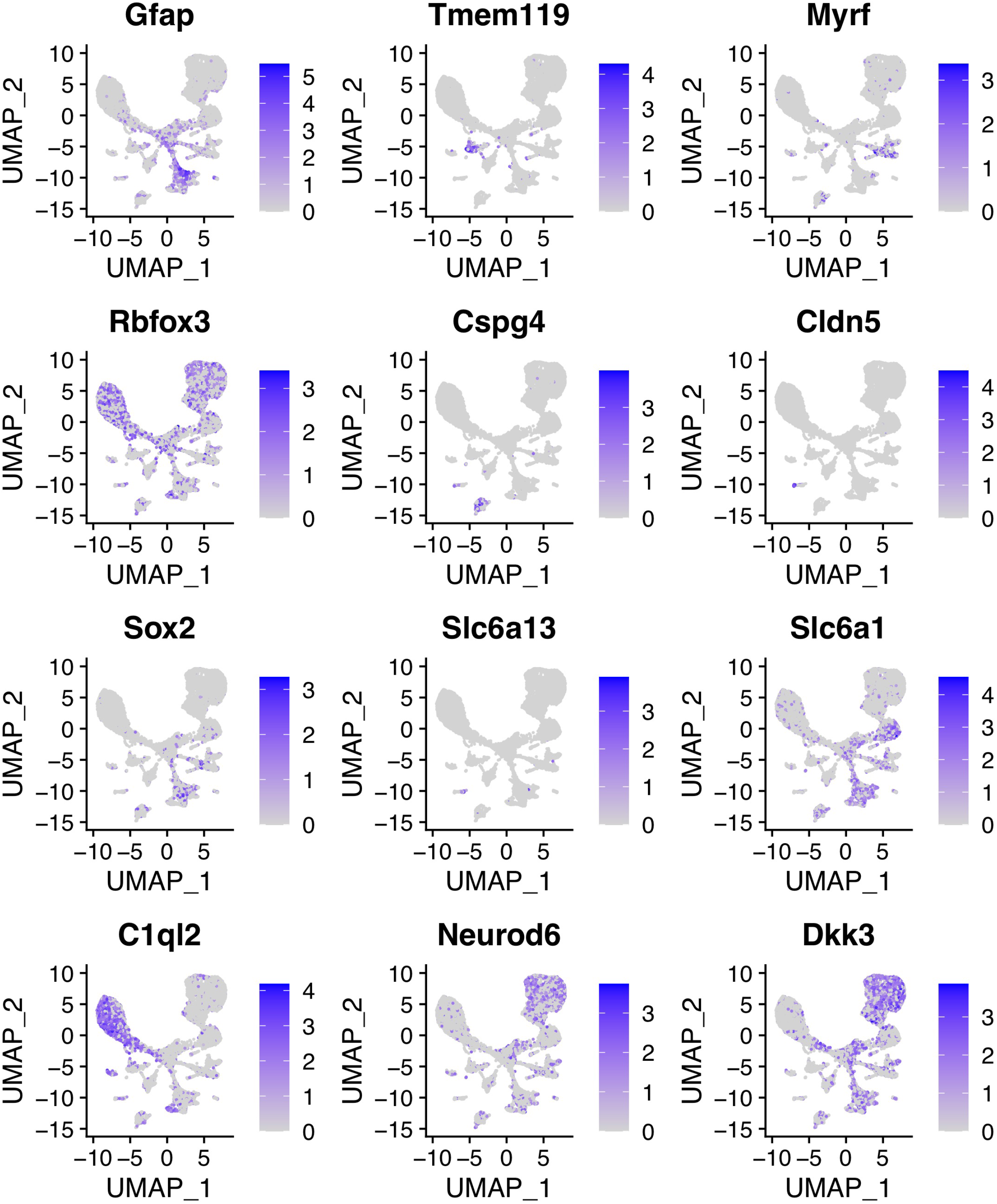
Marker gene expression for mouse hippocampal cells. Gfap = astrocytes, Tmem119 = microglia, Myrf = oligodendrocytes, Rbfox3 = neurons, Cspg4 = OPCs, Cldn5 = ependymal, Slc6a13 = vascular, Slc6a1 = interneurons, c1ql2 = DG neurons, Neurod6/Dkk3 = CA neurons.

## References

1. Burns KH, Boeke JD. Human transposon tectonics. Cell. 2012;149(4):740–752. doi:10.1016/j.cell.2012.04.019

2. Swergold GD. Identification, characterization, and cell specificity of a human LINE-1 promoter. Mol Cell Biol. 1990;10(12):6718–6729. doi:10.1128/mcb.10.12.6718-6729.1990

3. Wei W, Gilbert N, Ooi SL, et al. Human L1 Retrotransposition: cis Preference versus trans Complementation. Mol Cell Biol. 2001;21(4):1429–1439. doi:10.1128/MCB.21.4.1429-1439.2001

4. Martin SL. The ORF1 protein encoded by LINE-1: structure and function during L1 retrotransposition. J Biomed Biotechnol. 2006;2006(1):45621. doi:10.1155/JBB/2006/45621

5. Feng Q, Moran JV, Kazazian HH, Boeke JD. Human L1 retrotransposon encodes a conserved endonuclease required for retrotransposition. Cell. 1996;87(5):905–916.

6. Mathias SL, Scott AF, Kazazian HH, Boeke JD, Gabriel A. Reverse transcriptase encoded by a human transposable element. Science. 1991;254(5039):1808–1810. doi:10.1126/science.1722352

7. Mita P, Wudzinska A, Sun X, et al. LINE-1 protein localization and functional dynamics during the cell cycle. eLife. doi:10.7554/eLife.30058

8. Cost GJ, Feng Q, Jacquier A, Boeke JD. Human L1 element target-primed reverse transcription in vitro. EMBO J. 2002;21(21):5899–5910.

9. Hancks DC, Kazazian HH. Roles for retrotransposon insertions in human disease. Mob DNA. 2016;7:9. doi:10.1186/s13100-016-0065-9

10. Miki Y, Nishisho I, Horii A, et al. Disruption of the APC gene by a retrotransposal insertion of L1 sequence in a colon cancer. Cancer Res. 1992;52(3):643–645.

11. Scott EC, Gardner EJ, Masood A, Chuang NT, Vertino PM, Devine SE. A hot L1 retrotransposon evades somatic repression and initiates human colorectal cancer. Genome Res. 2016;26(6):745–755. doi:10.1101/gr.201814.115

12. Cajuso T, Sulo P, Tanskanen T, et al. Retrotransposon insertions can initiate colorectal cancer and are associated with poor survival. Nat Commun. 2019;10(1):4022. doi:10.1038/s41467-019-11770-0

13. Gasior SL, Wakeman TP, Xu B, Deininger PL. The Human LINE-1 Retrotransposon Creates DNA Double-strand Breaks. J Mol Biol. 2006;357(5):1383–1393. doi:10.1016/j.jmb.2006.01.089

14. Ardeljan D, Steranka JP, Liu C, et al. Cell fitness screens reveal a conflict between LINE-1 retrotransposition and DNA replication. Nat Struct Mol Biol. 2020;27(2):168–178. doi:10.1038/s41594-020-0372-1

15. Thomas CA, Tejwani L, Trujillo CA, et al. Modeling of TREX1-Dependent Autoimmune Disease using Human Stem Cells Highlights L1 Accumulation as a Source of Neuroinflammation. Cell Stem Cell. 2017;21(3):319–331.e8. doi:10.1016/j.stem.2017.07.009

16. Cecco MD, Ito T, Petrashen AP, et al. L1 drives IFN in senescent cells and promotes age-associated inflammation. Nature. 2019;566(7742):73. doi:10.1038/s41586-018-0784-9

17. Tunbak H, Enriquez-Gasca R, Tie CHC, et al. The HUSH complex is a gatekeeper of type I interferon through epigenetic regulation of LINE-1s. Nat Commun. 2020;11(1):5387. doi:10.1038/s41467-020-19170-5

18. Tang Z, Steranka JP, Ma S, et al. Human transposon insertion profiling: Analysis,visualization and identification of somatic LINE-1 insertions in ovarian cancer. Proc Natl Acad Sci U S A. 2017;114(5):E733–E740. doi:10.1073/pnas.1619797114

19. Rodić N, Steranka JP, Makohon-Moore A, et al. Retrotransposon insertions in the clonal evolution of pancreatic ductal adenocarcinoma. Nat Med. 2015;21(9):1060–1064. doi:10.1038/nm.3919

20. Tubio JMC, Li Y, Ju YS, et al. Mobile DNA in cancer. Extensive transduction of nonrepetitive DNA mediated by L1 retrotransposition in cancer genomes. Science. 2014;345(6196): 1251343. doi:10.1126/science.1251343

21. Rodriguez-Martin B, Alvarez EG, Baez-Ortega A, et al. Pan-cancer analysis of whole genomes identifies driver rearrangements promoted by LINE-1 retrotransposition. Nat Genet. 2020;52(3):306–319. doi:10.1038/s41588-019-0562-0

22. Helman E, Lawrence MS, Stewart C, Sougnez C, Getz G, Meyerson M. Somatic retrotransposition in human cancer revealed by whole-genome and exome sequencing. Genome Res. 2014;24(7):1053–1063. doi:10.1101/gr.163659.113

23. Jung H, Choi JK, Lee EA. Immune signatures correlate with L1 retrotransposition in gastrointestinal cancers. Genome Res. 2018;28(8):1136–1146. doi:10.1101/gr.231837.117

24. Gardner EJ, Lam VK, Harris DN, et al. The Mobile Element Locator Tool (MELT):Population-scale mobile element discovery and biology. Genome Res. Published online August 30, 2017:gr.218032.116. doi:10.1101/gr.218032.116

25. Skowronski J, Singer MF. Expression of a cytoplasmic LINE-1 transcript is regulated in a human teratocarcinoma cell line. Proc Natl Acad Sci U S A. 1985;82(18):6050–6054. doi:10.1073/pnas.82.18.6050

26. Martin SL. Ribonucleoprotein particles with LINE-1 RNA in mouse embryonal carcinoma cells. Mol Cell Biol. 1991;11(9):4804–4807. doi:10.1128/mcb.11.9.4804-4807.1991

27. Hickey DA. Selfish DNA: a sexually-transmitted nuclear parasite. Genetics. 1982;101(3-4):519–531. doi:10.1093/geneticsI101.3-4.519

28. Packer AI, Manova K, Bachvarova RF. A discrete LINE-1 transcript in mouse blastocysts. Dev Biol. 1993;157(1):281–283. doi:10.1006/dbio.1993.1133

29. Muotri AR, Chu VT, Marchetto MCN, Deng W, Moran JV, Gage FH. Somatic mosaicism in neuronal precursor cells mediated by L1 retrotransposition. Nature. 2005;435(7044):903–910. doi:10.1038/nature03663

30. Bestor TH. Cytosine methylation mediates sexual conflict. Trends Genet TIG. 2003;19(4):185–190. doi:10.1016/S0168-9525(03)00049-0

31. Kano H, Godoy I, Courtney C, et al. L1 retrotransposition occurs mainly in embryogenesis and creates somatic mosaicism. Genes Dev. 2009;23(11):1303–1312. doi:10.1101/gad.1803909

32. Simon M, Van Meter M, Ablaeva J, et al. LINE1 Derepression in Aged Wild-Type and SIRT6-Deficient Mice Drives Inflammation. Cell Metab. 2019;29(4):871–885.e5. doi:10.1016/j.cmet.2019.02.014

33. De Cecco M, Criscione SW, Peterson AL, Neretti N, Sedivy JM, Kreiling JA. Transposable elements become active and mobile in the genomes of aging mammalian somatic tissues. Aging. 2013;5(12):867–883.

34. Sun X, Wang X, Tang Z, et al. Transcription factor profiling reveals molecular choreography and key regulators of human retrotransposon expression. Proc Natl Acad Sci U S A. 2018;115(24):E5526–E5535. doi:10.1073/pnas.1722565115

35. Laughney AM, Hu J, Campbell NR, et al. Regenerative lineages and immune-mediated pruning in lung cancer metastasis. Nat Med. 2020;26(2):259–269. doi:10.1038/s41591-019-0750-6

36. Rodić N, Sharma R, Sharma R, et al. Long Interspersed Element-1 Protein Expression Is a Hallmark of Many Human Cancers. Am J Pathol. 2014;184(5):1280–1286. doi:10.1016/j.ajpath.2014.01.007

37. McKerrow W, Fenyö D. L1EM: a tool for accurate locus specific LINE-1 RNA quantification. Bioinforma Oxf Engl. 2020;36(4):1167–1173. doi:10.1093/bioinformatics/btz724

38. Deininger P, Morales ME, White TB, et al. A comprehensive approach to expression of L1 loci. Nucleic Acids Res. 2017;45(5):e31. doi:10.1093/nar/gkw1067

39. vdj_v1_hs_nsclc_5gex -Datasets-Single Cell Immune Profiling -Official 10x Genomics Support. Accessed October 22, 2020. https://support.10xgenomics.com/single-cell-vdj/datasets/2.2.0/vdj_v1_hs_nsclc_5gex

40. He S' Wang LH, Liu Y, et al. Single-cell transcriptome profiling of an adult human cell atlas of 15 major organs. Genome Biol. 2020;21(1):294. doi:10.1186/s13059-020-02210-0

41. Achanta P, Steranka JP, Tang Z, et al. Somatic retrotransposition is infrequent in glioblastomas. Mob DNA. 2016;7. doi:10.1186/s13100-016-0077-5

42. Carreira PE, Ewing AD, Li G, et al. Evidence for L1-associated DNA rearrangements and negligible L1 retrotransposition in glioblastoma multiforme. Mob DNA. 2016;7:21. doi:10.1186/s13100-016-0076-6

43. Wang D, Eraslan B, Wieland T, et al. A deep proteome and transcriptome abundance atlas of 29 healthy human tissues. Mol Syst Biol. 2019;15(2):e8503. doi:10.15252/msb.20188503

44. Satpathy S, Krug K, Beltran PMJ, et al. A proteogenomic portrait of lung squamous cell carcinoma. Cell. 2021;184(16):4348–4371.e40. doi:10.1016/j.cell.2021.07.016

45. Sookdeo A, Hepp CM, McClure MA, Boissinot S. Revisiting the evolution of mouse LINE-1in the genomic era. Mob DNA. 2013;4:3. doi:10.1186/1759-8753-4-3

46. McKerrow W, Wang X, Mita P, et al. LINE-1 expression in cancer correlates with DNA damage response, copy number variation, and cell cycle progression. bioRxiv. Published online June 28, 2020:2020.06.26.174052. doi:10.1101/2020.06.26.174052

47. Ardeljan D, Wang X, Oghbaie M, et al. LINE-1 ORF2p expression is nearly imperceptible in human cancers. Mob DNA. 2020;11:1. doi:10.1186/s13100-019-0191-2

48. Krug L, Chatterjee N, Borges-Monroy R, et al. Retrotransposon activation contributes to neurodegeneration in a Drosophila TDP-43 model of ALS. PLOS Genet. 2017;13(3):e1006635. doi:10.1371/journal.pgen.1006635

49. Tam OH, Rozhkov NV, Shaw R, et al. Postmortem Cortex Samples Identify Distinct Molecular Subtypes of ALS: Retrotransposon Activation, Oxidative Stress, and Activated Glia. Cell Rep. 2019;29(5):1164–1177.e5. doi:10.1016/j.celrep.2019.09.066

50. Guo C, Jeong HH, Hsieh YC, et al. Tau Activates Transposable Elements in Alzheimer's Disease. Cell Rep. 2018;23(10):2874–2880. doi:10.1016/j.celrep.2018.05.004

51. Carter V, LaCava J, Taylor MS, et al. High prevalence and disease correlation of autoantibodies against p40 encoded by long interspersed nuclear elements (LINE-1) in systemic lupus erythematosus. Arthritis Rheumatol Hoboken NJ. Published online July 24, 2019. doi:10.1002/art.41054

52. Crow MK. Long interspersed nuclear elements (LINE-1): potential triggers of systemic autoimmune disease. Autoimmunity. 2010;43(1):7–16. doi:10.3109/08916930903374865

53. Pizarro JG, Cristofari G. Post-Transcriptional Control of LINE-1 Retrotransposition by Cellular Host Factors in Somatic Cells. Front Cell Dev Biol. 2016;4:14. doi:10.3389/fcell.2016.00014

54. Wood J, Helfand S. Chromatin structure and transposable elements in organismal aging. Front Genet. 2013;4:274. doi:10.3389/fgene.2013.00274

55. Gorbunova V, Seluanov A, Mita P, et al. The role of retrotransposable elements in ageing and age-associated diseases. Nature. 2021;596(7870):43–53. doi:10.1038/s41586-021-03542-y

56. Bao W, Kojima KK, Kohany O. Repbase Update, a database of repetitive elements in eukaryotic genomes. Mob DNA. 2015;6(1):11. doi:10.1186/s13100-015-0041-9

57. Javahery R, Khachi A, Lo K, Zenzie-Gregory B, Smale ST. DNA sequence requirements for transcriptional initiator activity in mammalian cells. Mol Cell Biol. 1994;14(1):116–127. doi:10.1128/mcb.14.1.116-127.1994

58. Smale ST, Kadonaga JT. The RNA polymerase II core promoter. Annu Rev Biochem. 2003;72:449–479. doi:10.1146/annurev.biochem.72.121801.161520

59. Cembrowski MS, Wang L, Sugino K, Shields BC, Spruston N. Hipposeq: a comprehensive RNA-seq database of gene expression in hippocampal principal neurons. Marder E, ed. eLife. 2016;5:e14997. doi:10.7554/eLife.14997

60. Frazzette N, Khodadadi-Jamayran A, Doudican N, et al. Decreased cytotoxic T cells and TCR clonality in organ transplant recipients with squamous cell carcinoma. Npj Precis Oncol. 2020;4(1):1–8. doi:10.1038/s41698-020-0119-9

